# Transcriptomic analysis of diverse organisms reveals the rareness of genetic assimilation in environmental adaptations

**DOI:** 10.1101/2023.02.18.529054

**Authors:** Piaopiao Chen, Jianzhi Zhang

## Abstract

Genetic assimilation is the evolutionary process by which an environmentally induced phenotype becomes genetically encoded and constitutive. Genetic assimilation has been proposed as the concluding step in the plasticity-first model of environmental adaptation and has been observed in multiple species, but its prevalence has not been systematically investigated. By analyzing transcriptomic data collected upon reciprocal transplant, we address this question in the experimental evolution, domestication, or natural evolution of a bacterial, a fungal, a plant, and four animal species. We find that genetic assimilation of environment-induced gene expression is the exception rather than the rule and that substantially more genes retain than lose their expression plasticity upon organismal adaptations to new environments. The probability of genetic assimilation varies among genes and decreases with the number of transcription factors controlling the gene and the expression level of the gene, supporting the hypothesis that genetic assimilation results primarily from passive losses of gene regulations that are not mutationally robust. Therefore, at the level of gene expression, our findings argue against the purported importance of genetic assimilation to environmental adaptation.

## INTRODUCTION

Phenotypic plasticity, the ability of a genotype to produce different phenotypes when exposed to different environments, appears to be an intrinsic property of all forms of life [1]. However, the origin, maintenance, and evolution of phenotypic plasticity are not well understood and the role of phenotypic plasticity in environmental adaptation is controversial [2–4]. Specifically, the extended evolutionary synthesis maintains that phenotypic plasticity plays a pioneering role in environmental adaptation and that this crucial contribution of phenotypic plasticity is underappreciated by the modern synthesis, the reigning theory of evolutionary biology since the 1930s and 1940s [5–12]. According to the prevailing model of environmental adaptation advocated by the extended evolutionary synthesis—“plasticity-first” [9, 13], when the environment changes, the reaction norm (i.e., the range of phenotypes expressed by a single genotype across environments) allows the production of a novel phenotype that is partially or fully adapted to the new environment. If natural selection keeps operating only in the new environment, the adaptive phenotype becomes genetically assimilated or encoded such that it is expressed not only in the new environment but also in the old environment. In other words, the reaction norm is modified and the phenotypic plasticity is lost upon the organismal adaptation to the new environment. Here, the evolutionary process by which an environmentally induced phenotype becomes genetically encoded and constitutive is known as genetic assimilation [14–16]. This concept was proposed by C. H. Waddington [17] and is best illustrated by a pair of artificial selection experiments he conducted in the 1950s on fruit flies. In the first experiment [18], Waddington induced the *crossveinless* wing phenotype with a 4-hr heat shock at 40°C at 17 to 23 hrs after puparium formation. He then selected for this phenotype upon heat shock for 23 generations; eventually the phenotype appeared even without heat shock. In the second experiment [19], Waddington observed that a fraction of flies developed a second thorax when he exposed fly embryos to ether vapor. He selected for this *bithorax* phenotype with the ether treatment for 19 generations, after which some flies developed *bithorax* without the ether treatment. In both cases, a phenotype that is initially expressed only in a new environment becomes constitutive after generations of selection for the phenotype in the new environment. The concept of genetic assimilation was later expanded to genetic accommodation, which not only allows a decrease in phenotypic plasticity upon environmental adaptation but also an increase (i.e., the phenotype becomes more sensitive to the environment) [9]. We focus on genetic assimilation here because genetic accommodation appears too flexible to be a specific scientific hypothesis.

Adaptive plasticity and genetic assimilation (or the broader genetic accommodation) are considered two key components of the plasticity-first model of environmental adaptation [10]. Although both adaptive plasticity and genetic assimilation are empirically supported in some cases [10, 16, 20–27], most past studies analyzed a very small number of phenotypic traits per study despite that many traits evolve during typical environmental adaptations. Realizing this problem, a number of authors have recently turned to transcriptomic data because gene expression changes are widely thought to play important roles in phenotypic plasticity and genetic adaptation [28] and because the expression level of every gene in the transcriptome can be considered a trait (albeit not necessarily independent from other such traits), allowing a general and unbiased evaluation of the plasticity-first model for a large group of traits. Overall, these studies found most plastic expression changes to be maladaptive (i.e., subsequent genetic expression changes reverse the plastic changes) instead of adaptive, casting doubt on the plasticity-first model [29–32]. Nevertheless, even when most plastic gene expression changes are maladaptive, one could ask whether those few adaptive plastic changes become genetically assimilated upon environmental adaptations. This question was not previously addressed probably because differentiating adaptive from maladaptive plasticity requires only one transplant experiment while testing genetic assimilation requires reciprocal transplants (see Results) that are less commonly performed.

In the present work, we collect all available transcriptome data from reciprocal transplant experiments that are amenable to the test of genetic assimilation. These datasets are associated with the experimental evolution, domestication, or natural evolution of seven diverse species. Analyzing these data allows addressing two questions in the context of gene expression plasticity and evolution. First, is genetic assimilation of adaptive plasticity common in environmental adaptations? Second, what features are associated with genetic assimilation? The second question is relevant to the understanding of the process of genetic assimilation, which in turn sheds light on the role of genetic assimilation in environmental adaptation. In addition to testing the genetic assimilation hypothesis as part of the plasticity-first model, our study generates perhaps the first general picture of the evolution of phenotypic plasticity for a very large group of traits during the environmental adaptations of diverse organisms.

## RESULTS

### Experimental design

To test genetic assimilation in the adaptive evolution of a population to a new environment, we need access to the ancestral genotype that is adapted to the ancestral environment and the derived genotype that is adapted to the new environment. Let us use O (for original) and A (for adapted) to denote the states of the above two genotypes in their respective environments (**Fig. 1a**). In addition, let us use P (for plastic) to denote the state of the ancestral genotype in the new environment and B (for back) to denote the state of the derived genotype when it is moved back to the ancestral environment (**Fig. 1a**). Here, P and B can only be assessed after reciprocally transplanting the two genotypes from their native environments to the alternative environments (**Fig. 1a**). Let us use *E*_O_, *E*_P_, *E*_A_, and *E*_B_ to denote the expression level of a gene of interest at the state of O, P, A, and B, respectively. As is clear from Waddington’s experiments [18, 19], a prerequisite for genetic assimilation is that *E*_O_ ≠ *E*_A_, which limits our test of genetic assimilation to differentially expressed genes (DEGs) between O and A (**Fig. 1b**). Furthermore, because the genetic assimilation hypothesis is concerned with adaptive plasticity and because Waddington selected for the phenotype at state P in the new environment, we further limit our analysis to genes whose *E*_P_ is not significantly different from *E*_A_ but is significantly different from *E*_O_ (**Fig. 1c**), allowing testing the genetic assimilation hypothesis under a condition similar to Waddington’s experiments. We refer to DEGs with *E*_P_ not significantly different from *E*_A_ but significantly different from *E*_O_ plasticity-sufficient genes (PSGs), not because the gene expression is necessarily unaffected by genetic changes in evolution but because plasticity alone is sufficient to create state A from state O for these genes (**Fig. 1c**). While our investigation focused on PSGs for the reason explained, we also analyzed other genes and presented the results in supplementary materials (see Discussion).

**Fig. 1.**
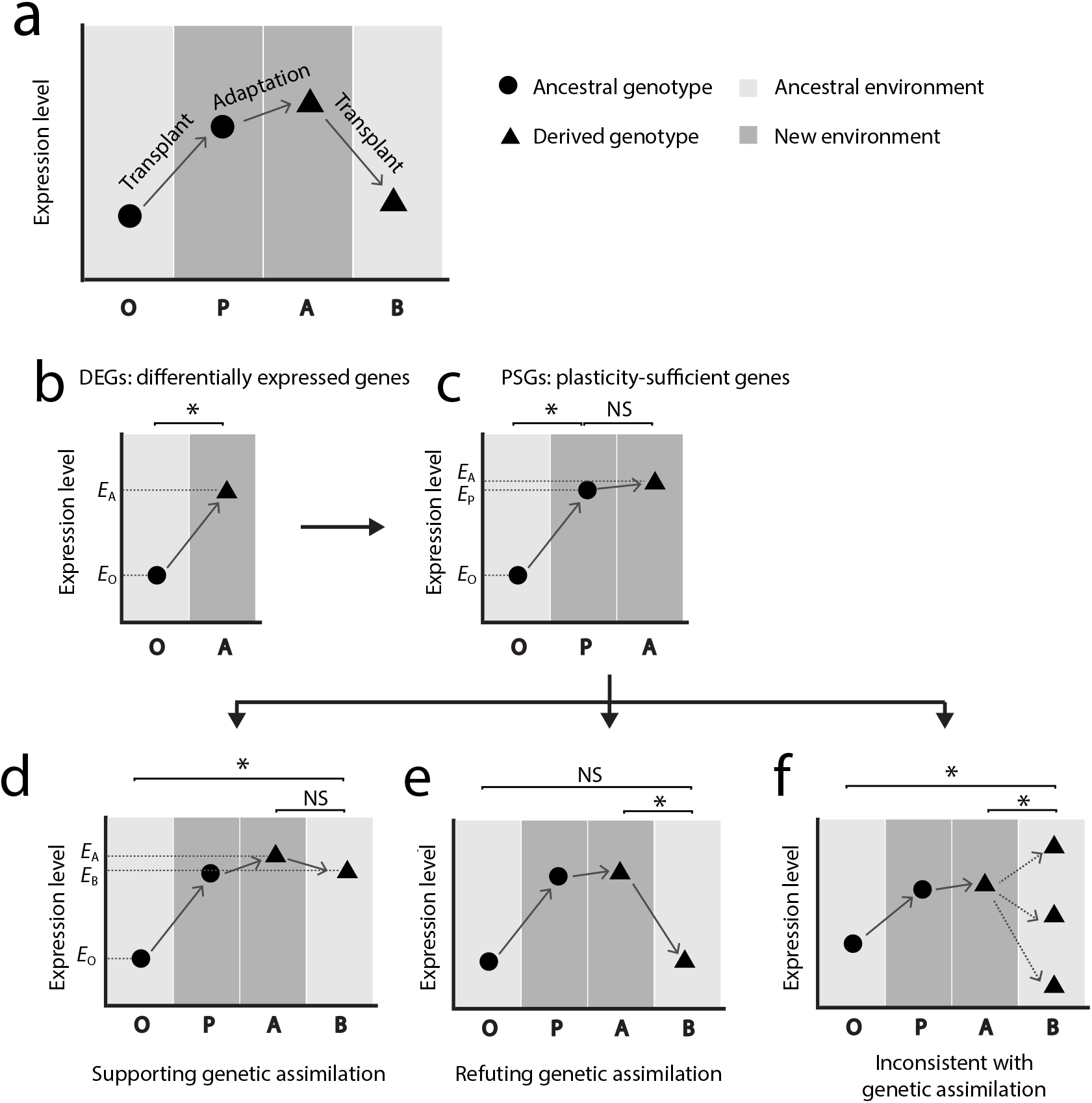
Experimental design for testing genetic assimilation of gene expression in environmental adaptation. **a**, Phenotyping the ancestral genotype that is adapted to the ancestral environment in the ancestral environment (state O) as well as in the new environment (state P), and phenotyping the derived genotype that is adapted to the new environment in the new environment (state A) as well as in the ancestral environment (state B). Reciprocal transplant is required to assess the phenotype of each genotype in its non-native environment. **b**, Identifying differentially expressed genes (DEGs), which have significantly different expression levels in states O and A. **c**, Identifying plasticity-sufficient genes (PSGs), which are DEGs with *E*_P_ not significantly different from *E*_A_ but significantly different from *E*_O_. **d**, Identifying PSGs that support genetic assimilation, which have *E*_B_ significantly different from *E*_O_ but not significantly different from *E*_A_. **e**, Identifying PSGs that refute genetic assimilation, which have *E*_B_ significantly different from *E*_A_ but not significantly different from *E*_O_. **f**, Additional PSGs that are inconsistent with genetic assimilation. These genes have *E*_B_ significantly different from both *E*_A_ and *E*_O_. Note that PSGs with *E*_B_ significantly different from neither *E*_A_ nor *E*_O_ are not informative for testing genetic assimilation so are not shown. *, expression levels are significantly different between the two states compared; NS, expression levels are not significantly different between the two states compared.

For a PSG, if its *E*_B_ is not significantly different from *E*_A_ but is significantly different from *E*_O_ (**Fig. 1d**), genetic assimilation has happened. We thus consider such PSGs to be supportive of the genetic assimilation hypothesis. Because the plasticity-first model posits that genetic assimilation is common (at least for adaptive plasticity) [10], the proportion of PSGs supporting genetic assimilation serves as one indicator of the generality of the plasticity-first model. Nonetheless, the proportion depends on one significant statistical test result and one non-significant result, so could be influenced by data quality and replicate number. In other words, a low proportion could occur for statistical reasons. We thus examine the following situation as a comparison. If *E*_B_ is significantly different from *E*_A_ but not significantly different from *E*_O_, genetic assimilation has not happened, so the PSG refutes the genetic assimilation hypothesis (**Fig. 1e**). The above two situations are exactly opposite to each other, so their abundances can be directly compared. If both proportions are low, the data may lack the power for assessing genetic assimilation; otherwise, the data are informative. In addition to the above two situations, there are three situations that are also inconsistent with genetic assimilation, all of which having *E*_B_ significantly different from both *E*_O_ and *E*_A_ (**Fig. 1f**). Because these three situations require two significant statistical test results whereas the first two situations require one significant test result and one non-significant test result, it would be unfair to compare their abundances. For this reason, we will focus on the first two situations in our analysis. Finally, if neither *E*_O_ nor *E*_A_ differs significantly from *E*_B_, the case may simply lack statistical power for evaluating genetic assimilation.

### *Escherichia coli* experimental evolution in 11 stressful environments

We searched the literature and found seven studies with available transcriptomic data that fit our experimental design for testing genetic assimilation. The largest in terms of the number of new environments examined was an *E. coli* experimental evolution study [33]. Specifically, *E. coli* was first adapted to a benign (ancestral) environment for approximately 90 generations, followed by evolution in one of 11 stressful environments for 350 to 800 generations. The adaptation of *E. coli* to each of the stressful environments was evident from the drastic increase in the population growth rate during the experimental evolution. Transcriptomic data were collected for the ancestral environment–adapted strain in the ancestral environment (O) and shortly after exposure to each stressful environment (P), as well as each stress-adapted strain in its respective stressful environment (A) and back in the ancestral environment (B). This dataset allowed testing genetic assimilation in 11 different environmental adaptations.

From each of the 11 adaptations, we first identified DEGs by requiring *E*_A_ to be significantly different from *E*_O_ at the false discovery rate (FDR) of 0.05 (**Fig. 2a**). We then identified PSGs from all DEGs by requiring that *E*_P_ is not significantly different from *E*_A_ but is significantly different from *E*_O_, at the FDR of 0.05 in each test (**Fig. 2a**). From all PSGs, we identified genes supporting genetic assimilation by requiring their *E*_B_ to be significantly different from *E*_O_ but not significantly different from *E*_A_, at the FDR of 0.05 in each test. The number of PSGs supporting genetic assimilation varies from 5 to 68 across the 11 environments, accounting for only 1.1–20.4% of PSGs or 0.5–8.9% of DEGs (**Fig. 2a**). When the results from the 11 environments are combined, 4.9% of PSGs or 3.0% of DEGs support genetic assimilation (**Fig. 2a**). A typical example supporting genetic assimilation is shown in **Fig. 2b**. Specifically, *nemA* expression rose significantly when *E. coli* cells were moved from the ancestral benign environment to the stressful methylglyoxal environment; upon *E. coli*’s adaptation to this environment, *nemA* became similarly highly expressed in the ancestral and stressful environments. Clearly, the original stress-induced high expression of *nemA* has been genetically assimilated upon *E. coli*’s adaptation to the stress.

**Fig. 2.**
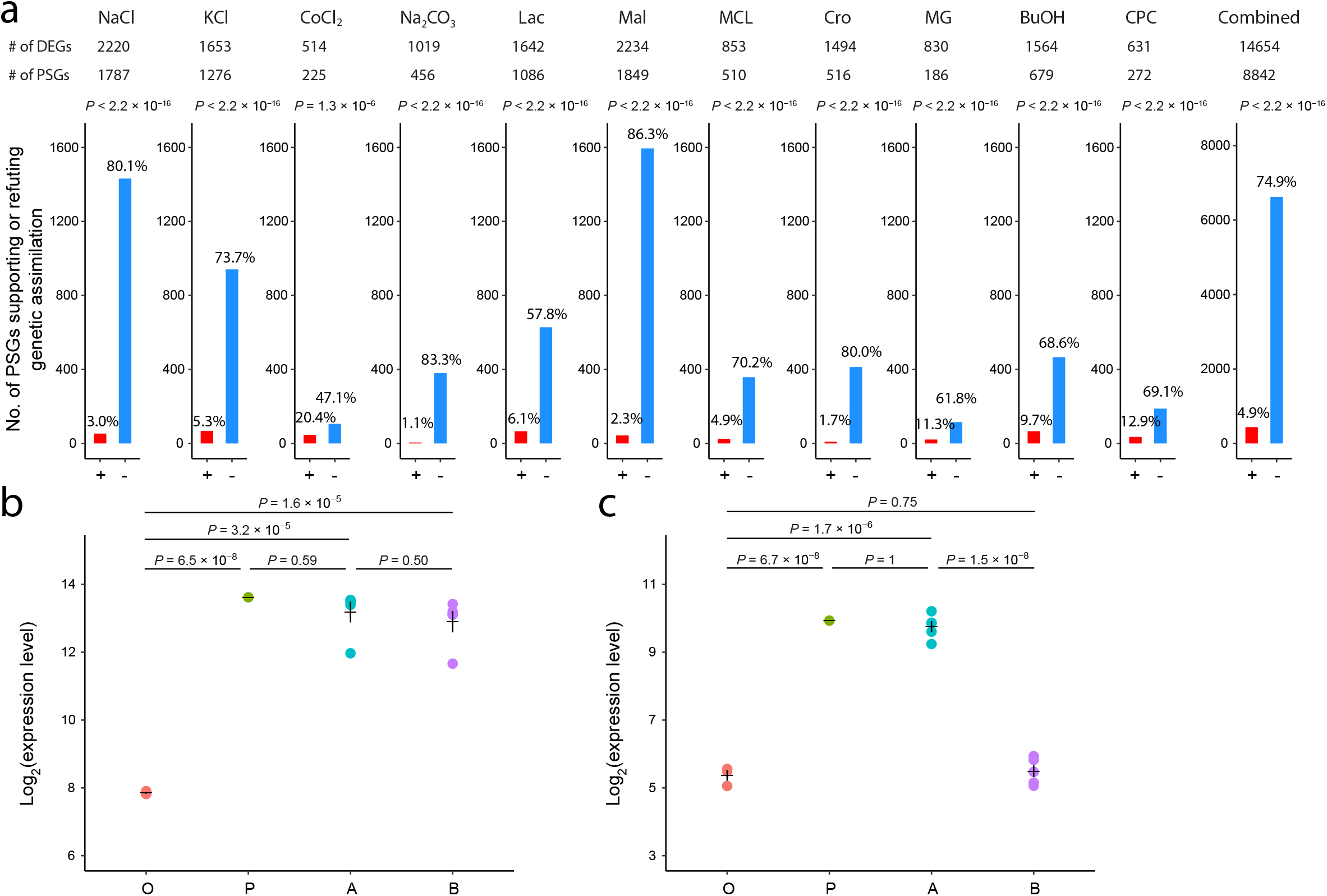
Genes supporting or refuting genetic assimilation in laboratory adaptations of *E. coli* to 11 different stressful environments. **a**, Numbers of DEGs, PSGs, and PSGs supporting (red bars) or refuting (blue bars) genetic assimilation in each of the 11 environments as well as for all environments combined. The number above each bar is the fraction of PSGs supporting or refuting genetic assimilation. + and - indicate supporting and refuting genetic assimilation, respectively. *P*-values are from two-tailed binomial tests of equal probabilities of supporting and refuting genetic assimilation. The 11 environments are indicated on the top of the panel and are detailed in Materials and Methods. **b**, Expression levels of a typical PSG (*nemA*) that supports genetic assimilation in the adaptation to the methylglyoxal medium. **c**, Expression levels of a typical PSG (*rygB*) that refutes genetic assimilation in the adaptation to the L-malate medium. In b-c, each dot represents the expression level of a replicate in state O, P, A, or B. Horizontal and vertical lines respectively show the mean and standard error of the expression level at each state. *P*-values are from moderated contrast *t*-test adjusted for multiple testing.

To exclude the possibility that the low numbers of genes supporting genetic assimilation are caused by statistical issues, we identified genes that refute the genetic assimilation hypothesis by requiring *E*_B_ to be significantly different from *E*_A_ but not significantly different from *E*_O_, at the FDR of 0.05 in each test. For example, the expression of *rygB* rose when *E. coli* was moved to the L-malate medium and stayed at approximately the same level upon *E. coli’s* adaptation to this medium (**Fig. 2c**). But when the malate-adapted *E. coli* was moved back to the ancestral environment, *rygB’s* expression decreased to the original level, indicating that genetic assimilation did not occur (**Fig. 2c**). The number of genes refuting genetic assimilation varies from 106 to 1,595 across the 11 environments, accounting for 47.1–86.3% of PSGs or 13.9–71.4% of DEGs. Combining the results from the 11 environments, we found 75% of PSGs or 45% of DEGs to refute genetic assimilation; these percentages are about 15 times those supporting genetic assimilation. To avoid the potential statistical bias resulting from the sample size difference in measuring *E*_0_ (*n* = 3 replicates) and *E*_A_ (*n* = 5 replicates), we randomly picked three from the five replicates in *E*_A_ measurement. We performed all ten possible downsamplings in each environment and used the mean of the ten FDR values as the final FDR for each gene. Again, the number of genes refuting genetic assimilation is overall 12 times that supporting genetic assimilation (**Fig. S1**). Clearly, genetic assimilation was the exception rather than the rule in the adaptations of *E. coli* to the 11 stresses. However, due to the difficulty in measuring the fitness effects of gene expression changes, the relative contributions of genes supporting and refuting genetic assimilation to environmental adaptations are unknown (see Discussion).

### Yeast experimental evolution in two stressful environments

Dhar *et al*. [34] performed experimental evolution of the baker’s yeast *Saccharomyces cerevisiae* that had been adapted to a benign environment in a high-salt (NaCl) medium or a medium with oxidative stress (H_2_O_2_) for 300 generations. They observed that yeast fitness in each stressful environment increased significantly upon the respective experimental evolution. Transcriptomic data representing states O, P, A, and B were collected for each of the two environmental adaptations. We identified from the transcriptome data 2,108 and 1,156 PSGs in salt and oxidative stress, respectively (**Fig. 3a**). Strikingly, we found no gene supporting genetic assimilation, while detecting 2,104 genes (99.8% of PSGs) and 1,141 genes (98.7% of PSGs) refuting genetic assimilation in the two environments, respectively (**Fig. 3a**).

**Fig. 3.**
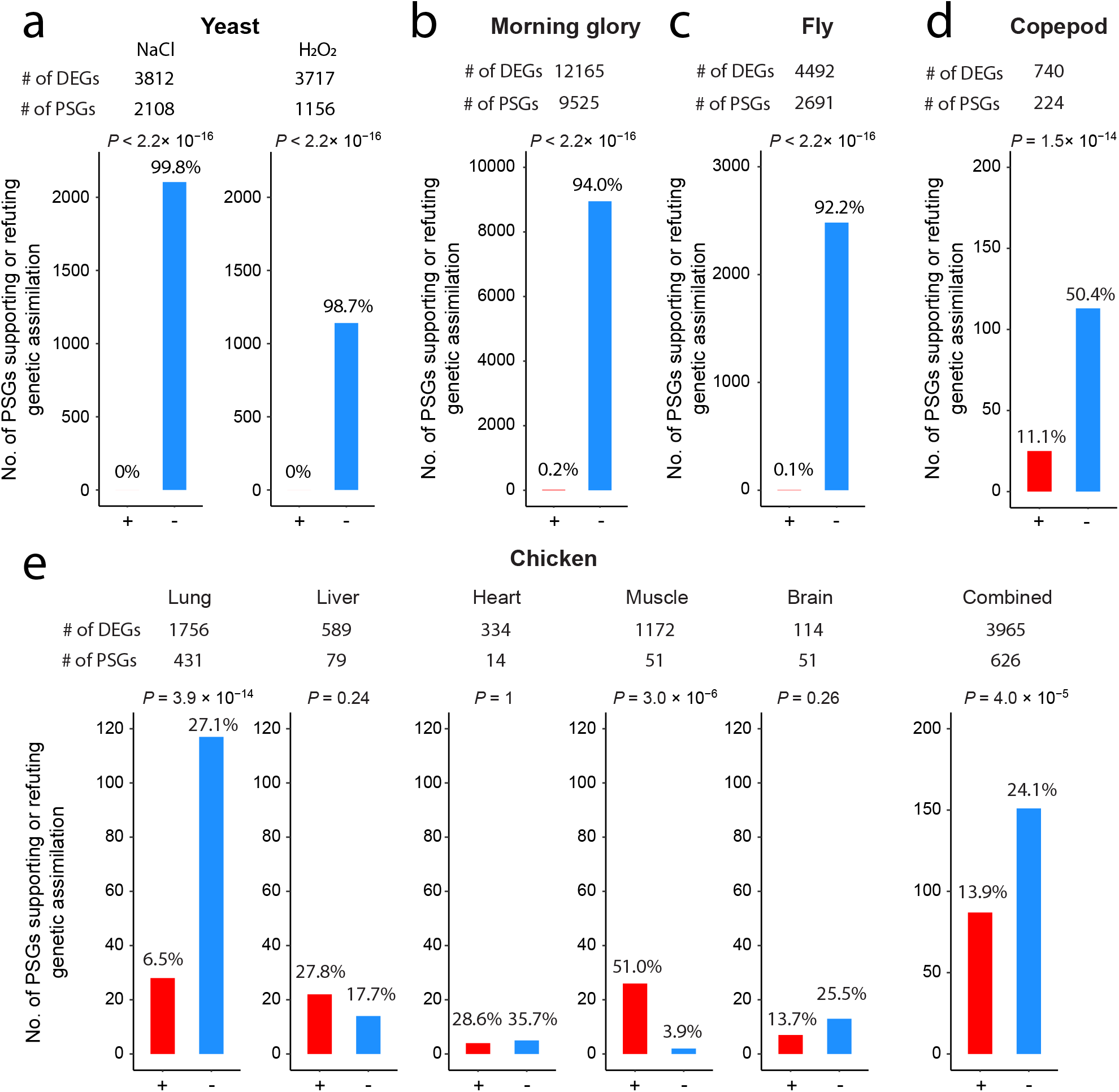
Genes supporting or refuting genetic assimilation in five other species studied. **a-e**, Numbers of DEGs, PSGs, and PSGs supporting (red bars) or refuting (blue bars) genetic assimilation in yeast adaptations to two separate stresses (**a**), morning glory’s evolution of herbicide resistance (**b**), fruit fly’s adaptation to a hot environment (**c**), copepod’s adaptation to a stressful environment (**d**), and Tibetan chicken’s adaptation to the highland (**e**). The number on the top of each bar is the fraction of PSGs supporting or refuting genetic assimilation. + and - indicate supporting and refuting genetic assimilation, respectively. *P*-values are from two-tailed binomial tests of equal probabilities of supporting and refuting genetic assimilation. In e, the results from each of the five tissues profiled as well as those for all tissues combined are shown.

### Selection of herbicide-resistant morning glories

Van Etten *et al*. [35] grew the common morning glory *Ipomoea-purpurea* in the herbicide (glyphosate) environment for three generations (~126 days) to obtain herbicide-resistant lines, and in the ancestral environment without the herbicide for the same number of generations as a control. Leaf tissues were then sampled from both experimentally evolved lines and control lines, followed by measurement of their transcriptomes at two time points (8 and 32 hrs) post-herbicide application, as well as in the herbicide-free environment at the same time points. We used the transcriptomic data from the later time point for each comparison to ensure a full environmental induction of gene expression. We found that 17 genes (0.2% of PSGs) support genetic assimilation, while 8,949 genes (94.0% of PSGs) refute it (**Fig. 3b**). Qualitatively similar results were obtained from the earlier time point (31 and 4,792 genes, respectively). Hence, there was little genetic assimilation in the morning glory’s evolution of herbicide-resistance.

### Laboratory adaptation of fruit flies to a hot environment

Mallard *et al*. [36, 37] exposed a *Drosophila simulans* population to a hot environment that fluctuated between 18°C and 28°C and a cold environment that fluctuated between 10°C and 20°C, for a total of 64 (hot) and 39 (cold) generations, respectively. Transcriptomic data were collected from the hot-evolved population at high (23°C) and low (15°C) temperatures, respectively, as well as from the cold-evolved population at the same two temperatures. We considered the cold laboratory environment to mimic the ancestral environment for two reasons. First, the flies were collected in 2008 from Póvoa de Varzim, Portugal, where the temperature was close to the cold laboratory environment. Second, in both environments, the similarity in gene expression profile between the ancestral population and the cold-evolved population is substantially higher than that between the ancestral population and the hot-evolved population (**Fig. S2**). Consistent with an adaptive response, the hot-evolved population was fitter than the ancestral population in the hot environment. We found that, among 2,691 PSGs, only 3 (0.1%) genes support genetic assimilation, substantially fewer than the 2,482 (92.2%) genes refuting genetic assimilation (**Fig. 3c**). Hence, genetic assimilation was rare in the laboratory adaptation of fruit flies to a hot environment.

### Experimental evolution of copepods in a stressful environment

Copepods are small crustaceans found in nearly every freshwater and saltwater habitat. To study their potential response to future climate changes, Brennan *et al*. [38] experimentally evolved the marine copepod *Acartia tonsa* in an environment representing today’s oceanic pH and temperature as well as in a stressful environment with a lower pH and a higher temperature for 20 generations. The copepod in the stressful environment rapidly adapted by recovering egg production and hatching success. Each population was then split into two, one maintained in the same environment as the previous 20 generations for three more generations while the other transplanted to the alternative environment for three generations. We analyzed the transcriptomic data collected at the end of the 23 generations. Of 224 PSGs, 25 (11.1%) support genetic assimilation, much fewer than the 113 (50.4%) genes refuting genetic assimilation (**Fig. 3d**).

### Experimental and natural evolution of Trinidadian guppies

Ghalambor *et al*. [29] studied the natural evolution of the Trinidadian guppy *Poecilia reticulata* that had been adapted to a high-predation environment in a new low-predation environment, as well as two replicates of a one-year experimental evolution in which the guppies were artificially introduced from the high-predation environment to the low-predation environment. The authors found indications of adaptations of the guppies to the low-predation environment [29]. They then sequenced the brain transcriptomes of the guppies at states O, P, A, and B in each of the three parallel environmental adaptations. Interestingly, in the natural evolution case, no PSG was observed among 678 DEGs; consequently, no genetic assimilation could happen. In the two experimental evolution cases, 25 and 20 DEGs were respectively detected, but again no PSG was found. Hence, genetic assimilation did not occur at all in the guppies.

### Highland adaptation of Tibetan chickens

The Tibetan chicken, well adapted to the highland, is thought to be derived from the lowland (domesticated) chicken that were brought to the Tibetan Plateau by ~1200 years ago [39]. Ho *et al*. [39] respectively hatched and raised lowland chickens in the highland and Tibetan chickens in the lowland, followed by transcriptome profiling of five tissues (lung, liver, heart, muscle, and brain). In addition, the transcriptomic data for the same five tissues were collected from lowland chickens hatched and raised in the lowland as well as Tibetan chickens hatched and raised in the highland. From these data, we found 28 (6.5% of PSGs), 22 (27.8%), 4 (28.6%), 26 (51.0%), and 7 (13.7%) genes that support genetic assimilation in the lung, liver, heart, muscle, and brain, respectively (**Fig. 3e**). The number of genes supporting genetic assimilation is significantly lower than that refuting genetic assimilation in the lung, while the opposite is true in the muscle (**Fig. 3e**). The two numbers are not significantly different from each other for each of the other three tissues examined (**Fig. 3e**). Nonetheless, combined across the five tissues, the number of genes supporting genetic assimilation (87) is significantly smaller than that (151) refuting genetic assimilation (*P* = 4.0×10^−5^, two-tailed binomial test). Hence, despite the variation across tissues, genetic assimilation is overall relatively infrequent in the highland adaptation of Tibetan chickens.

### Probability of genetic assimilation varies among genes

While genetic assimilation is overall rare in the environmental adaptations of the seven diverse species studied here, we may still use the limited number of cases to investigate whether genetic assimilation is equally likely for different genes, and if the answer is no, what features impact the probability of genetic assimilation. To this end, we further analyzed the *E. coli* data because of the availability of 11 sets of genes with genetic assimilation and abundant biological knowledge of this model organism. We performed Gene Ontology (GO) analysis of all *E. coli* genes supporting genetic assimilation (against the background of all PSGs), and found significant enrichments with electron transport chain, cellular respiration, succinate dehydrogenase activity, and several metabolic processes (**Fig. S3**). In addition, we performed GO analysis in the other six species studied and found genes supporting genetic assimilation to be significantly enriched with the GO terms of developmental growth in morning glories, and extracellular space in chickens. These enriched GO terms may be related to the specific environmental adaptations of each species so do not overlap among species, suggesting that genes exhibiting genetic assimilation in expression are idiosyncratic to specific organisms and environments.

A total of 68 genes of *E. coli* show genetic assimilation in at least two of the 11 environmental adaptations and 10 of them exhibit genetic assimilation in at least four environmental adaptations (**Fig. 4a**). The 68 genes show up in 50 different pairs of environments, suggesting that this observation is not due to the potential similarity among some test environments. Given the large number of *E. coli* genes in the genome, this pattern suggests unequal probabilities of genetic assimilation across genes. Under the null hypothesis of an equal probability of genetic assimilation for all genes, the number of new environments in which a gene exhibits genetic assimilation should approximately follow a Poisson distribution, whose variance equals the mean. The actual distribution, however, has a much greater variance than the mean (*P* < 2.2×10^−16^, chi-squared test of the null hypothesis that the observed variance equals the mean), with overrepresentations of genes showing genetic assimilation in 0 and ≥ 2 environmental adaptations (**Fig. 4a**). To exclude the possibility that the observed overdispersion is entirely due to overlapping PSGs among the 11 environments, in each environment, instead of considering the actual genes exhibiting genetic assimilation, we randomly sampled the same number of genes from the PSGs and treated them as genetically assimilated genes. This process was repeated to obtain 10,000 fake sets of genetically assimilated genes from 11 environmental adaptations. Compared with the average of the 10,000 fake sets, the actual assimilated genes still show overrepresentations in 0 and ≥ 2 environmental adaptations (**Fig. 4b**), demonstrating a genuine variation in the probability of genetic assimilation across genes.

**Fig. 4.**
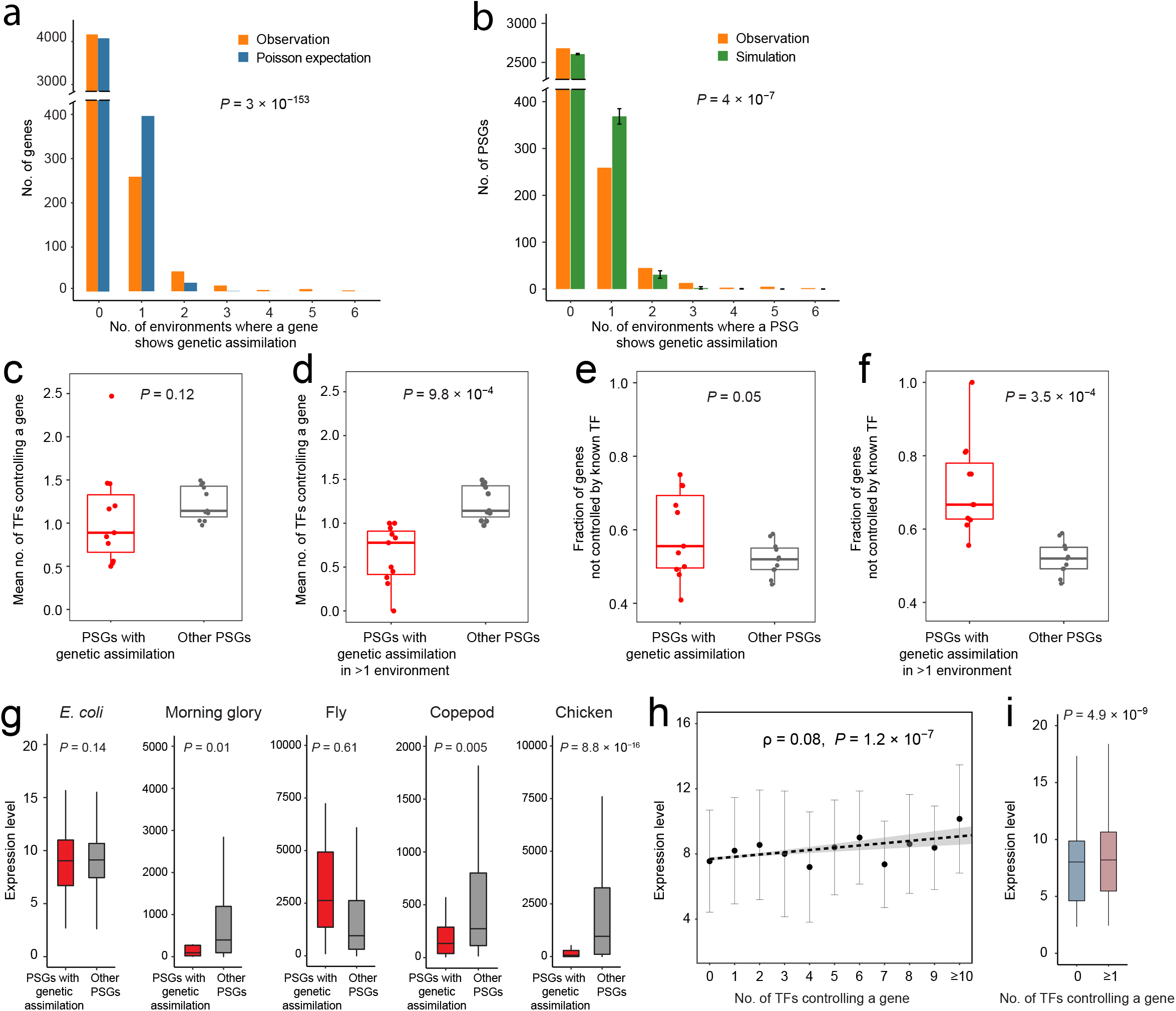
Characteristics of genes exhibiting genetic assimilation. **a**, Frequency distribution of the number of environments where a gene exhibits genetic assimilation in the *E. coli* data, compared with the Poisson distribution with the same mean. *P*-value is based on a one-tailed chi-squared test comparing the variance of the observed distribution with the corresponding Poisson variance. **b**, Frequency distribution of the number of environments where a PSG exhibits genetic assimilation in the *E. coli* data, compared with the corresponding distribution from the average of 10,000 simulations in which genes exhibiting genetic assimilation in each environment are randomly picked from the PSGs of the environment. Error bars indicate the 5th and 95th percentiles of the simulated data. *P*-value is from a one-tailed chi-squared test. **c**, Mean number of TFs controlling a gene for the group of genes exhibiting genetic assimilation or for the rest of PSGs. **d**, Mean number of TFs controlling a gene for the group of genes exhibiting genetic assimilation in at least two environments or for the rest of PSGs. **e**, Fraction of genes not controlled by any known TF for the group of genes exhibiting genetic assimilation or for the rest of PSGs. **f**, Fraction of genes not controlled by any known TF for the group of genes exhibiting genetic assimilation in at least two environments or for the rest of PSGs. In c-f, each dot represents one of the 11 environmental adaptations of *E. coli*. In the box plot, the lower and upper edges of the box represent the first and third quartiles, respectively, the horizontal line inside the box indicates the median, and the whiskers extend to the most extreme values inside inner fences (median ± 1.5× interquartile range). *P* values are from one-tailed paired *t*-test. **g**, Gene expression distribution in the ancestral environment for genes exhibiting genetic assimilation or the rest of PSGs. In *E. coli*, a gene is considered to exhibit genetic assimilation if it exhibits genetic assimilation in at least one of the 11 environmental adaptations. *P* values are from two-tailed Wilcoxon rank-sum tests. **h**, Number of TFs controlling an *E. coli* gene is positively correlated with its expression level in the ancestral environment. Dot and error bar respectively represent the mean and standard deviation of each category. The linear regression is shown by a dotted line. Spearman’s rank correlation (*ρ*) based on the unbinned data and its associated *P* value are presented. **i**, In the ancestral environment, expression levels of *E. coli* genes controlled by no known TF is significantly lower than those of genes controlled by at least one TF. *P*-value is from two-tailed Wilcoxon rank-sum test. In g-i, expressions of *E. coli* genes are normalized via the robust multi-array average (RMA) method and presented as log_2_-transformed values by RMA. Gene expressions in other organisms are presented as read counts normalized by the median of ratios method used by DESeq2 [55].

### Features of genes showing genetic assimilations

Our analysis of genetic assimilation focused on the mRNA levels of genes. Because the mRNA level of a gene is largely determined by its transcriptional level, which is mainly controlled by transcription factors (TFs), the genetic assimilation revealed here may be caused primarily by an evolutionary change in the TF-based control of gene transcription in the ancestral environment. Because TFs often work in a redundant fashion [40, 41], we reasoned that the expressions of genes controlled by fewer TFs are likely to be less robust to any changes in TF or TF-binding, so are more likely to exhibit genetic assimilation. To test this hypothesis in *E. coli* where the information of genes controlled by each TF is readily available, we computed the mean number of TFs controlling genes exhibiting genetic assimilation and the corresponding value for all other PSGs in each environmental adaptation; the former is smaller than the latter across the 11 environments, although the difference is not statistically significant (*P* = 0.12, one-tailed paired *t*-test; **Fig. 4c**). This difference becomes highly significant between genes exhibiting genetic assimilation in at least two environments and the rest of PSGs (*P* < 10 ^−3^; **Fig. 4d**). To further test the hypothesis, we computed the fraction of genes controlled by no known TFs among those with genetic assimilation and the corresponding fraction among other PSGs in each environmental adaptation; as expected, the former is significantly higher than the latter across the 11 environments (*P* = 0.05, **Fig. 4e**). Similarity, this disparity is much more significant between genes exhibiting genetic assimilation in at least two environmental adaptations and the remaining PSGs (*P* < 10^−3^; **Fig. 4f**).

We further examined whether the gene expression level in the ancestral environment has any effect on the probability of genetic assimilation. We observed in *E. coli* lower expressions for genes with genetic assimilation than other PSGs at face values, but the difference is not statistically significant (*P* = 0.14, two-tailed Wilcoxon rank-sum test; **Fig. 4g**). This trend is more pronounced and is significant in the morning glory, copepod, and chicken (**Fig. 4g**), is reversed but not statistically significant in the fly (**Fig. 4g**), and cannot be assessed in the yeast or guppy due to their lack of genes with genetic assimilation. This trend may be due to a positive correlation between the number of TFs controlling a gene and the expression level of the gene. Indeed, we were able to confirm this correlation in *E. coli* (**Fig. 4h**). Furthermore, genes controlled by no known TFs have significantly lower expressions than genes controlled by at least one TF (**Fig. 4i**). Hence, the effect of gene expression level on the probability of genetic assimilation is probably mediated by the number of TFs controlling the gene.

## DISCUSSION

Analyzing the transcriptome data from environmental adaptations of diverse species, we found that genetic assimilation of gene expression is generally rare. The median proportion of PSGs supporting genetic assimilation is only 2.6% for the six species studied (guppies had no PSG so are excluded here, and the overall proportion of PSGs supporting genetic assimilation is considered for species with data from multiple environmental adaptations). For comparison, the corresponding value refuting genetic assimilation is 83.6%. Clearly, genetic assimilation is the exception rather than the rule of gene expression plasticity evolution in environmental adaptations.

In our test of genetic assimilation, we followed Waddington’s experiments [18, 19] to consider only PSGs, for which *E*_P_ is not significantly different from *E*_A_. However, Waddington studied discrete traits while gene expression levels are quantitative traits. For a DEG, if *E*_O_ and *E*_P_ are significantly unequal and *E*_P_ is somewhere between *E*_O_ and *E*_A_, the difference between *E*_O_ and *E*_P_ can be considered adaptive plasticity even when *E*_P_ and *E*_A_ are significantly unequal. Because the genetic assimilation hypothesis is concerned with all adaptive plasticity, we could assess genetic assimilation of PSGs as well as DEGs whose *E*_P_ is significantly different from both *E*_A_ and *E*_O_ and is somewhere between *E*_O_ and *E*_A_ (referred to as genes with both plastic and genetic changes or PGGs). We refer to the union of PSGs and PGGs as adaptive plasticity genes (APGs). We observed 5.0% of APGs to support genetic assimilation and 74.5% of APGs to refute genetic assimilation by combining the results from the 11 environmental adaptations of *E. coli* (**Fig. S4**). Furthermore, 0%, 0.2%, 0.2%, 12.1%, and 13.9% of APGs support genetic assimilation in the yeast (combined from two environments), morning glory, fruit fly, copepod, and chicken (combined from five tissues), respectively (**Fig. S5**), while 99.2%, 94.1%, 93.9%, 45.6%, and 24.6% of APGs refute genetic assimilation, respectively. No APG was observed in any of the three environmental adaptations of the guppy. Qualitatively similar results were obtained when PGGs were analyzed separately, although PGGs are substantially fewer than PSGs (**Fig. S6**). Furthermore, the proportion of PGGs and that of PSGs supporting genetic assimilation are not significantly different in most species studied (*P* > 0.05 in *E. coli*, yeast, fly, copepod, and chicken; chi-squared test). Hence, extending the analysis from PSGs to PGGs or APGs does not alter our conclusion of the rareness of genetic assimilation.

More broadly, when comparing genes whose expressions exhibit significant plasticity in the ancestral but not derived genotype (*E*_O_ and *E*_P_ are significantly different but *E*_A_ and *E*_B_ are not significantly different; Group 1) and those whose expressions exhibit significant plasticity in the derived but not ancestral genotype (*E*_A_ and *E*_B_ are significantly different but *E*_O_ and *E*_P_ are not significantly different; Group 2), we found significantly fewer genes in Group 1 than in Group 2 for *E. coli*, yeast, morning glory, fly, guppy, and chicken, while the opposite is true in copepod (**Fig. S7**). Hence, there are both gains and losses of expression plasticity upon environmental adaptations and if anything, gains outnumber losses in most species, as was noted by authors of some of the datasets analyzed here [37, 42].

Waddington’s experiments showed that genetic assimilation can occur in about 20 generations [18, 19], but several theoretical studies suggested that genetic assimilation should generally take orders of magnitude more generations [43–45]. The number of generations in the environmental adaptations studied here varied from three for morning glories [35] to at least 2400 for Tibetan chickens [39], and no significant correlation across the six species (guppies are excluded due to the lack of PSGs) exists between the number of generations of evolution and the fraction of PSGs supporting genetic assimilation (*P* = 0.13). The same correlation is, however, significant, across the 11 environmental adaptations of *E. coli (r* = 0.64, *P* = 0.033). But the biological interpretation of this correlation is complicated by the fact that, in the *E. coli* experiment, the number of generations of evolution is positively correlated with the population growth rate, which is negatively correlated with the severity of the environmental stress. Hence, it is unclear whether the positive correlation means that genetic assimilation accrues with the evolutionary time during an environmental adaptation. In the same vein, the seven species and associated datasets vary in many intercorrelated features and to what extent these features impact the fraction of genes supporting genetic assimilation relative to that refuting genetic assimilation is unclear and should be investigated in the future when more data become available.

When a population is in a new environment, gene expression in the previous environment is no longer under selection. Hence, gene expression in the previous environment might gradually deviate from its original value, and a fraction of genes may by chance have their *E*_B_ similar to *E*_A_ and thus show genetic assimilation. The fraction of PSGs with genetic assimilation is predicted to be negatively correlated with the mutational robustness of *E*_B_. This prediction is supported by our finding in *E. coli* that the propensity for genetic assimilation varies among genes and that genes controlled by fewer TFs and with lower expressions are more prone to genetic assimilation. More generally, in addition to being impacted by the mutational robustness of *E*_B_, the evolution of *E*_B_ is likely influenced by selection in the new environment because of pleiotropy [46–49]. That is, mutations fixed in the new environment for their benefits in the new environment may affect *E*_B_. These mutations may or may not affect *E*_A_ and can occur anywhere in the genome, making their impacts on *E*_B_ difficult to predict. It is thus unclear why *E*_B_ would necessarily evolve towards *E*_A_ (i.e., genetic assimilation) given the initial presence of plasticity before the population encounters the new environment. Rather, if adapting to every new environment leads to genetic assimilation (i.e., loss of plasticity), there would be no adaptive plasticity that the plasticity-first model requires when another new environment arrives. It has been suggested that plasticity may be costly to maintain [50], leading to a selection for its loss [16]. However, the cost of plasticity appears to be small [51] or have limited evidence [52].

One important caveat in our study is that, strictly speaking, we do not know whether the difference between *E*_O_ and *E*_A_ for a gene contributes to the environmental adaptation or is merely a consequence of the adaptation and is on its own neutral or even deleterious. Nevertheless, because of the general difficulty in establishing the causality between a phenotypic change and a fitness change, especially in different environments, this caveat also applies to most genetic assimilation studies of other types of phenotypic traits such as morphological and physiological traits. For example, 13 of the 21 best cases supporting the plasticity-first model [10] lack evidence for an adaptive role of the trait concerned; even for those purported to have such evidence, the adaptiveness of the focal trait is not without ambiguity because of the difficulty of separating multiple traits in fitness assessment. In the context of plasticity of gene expression, the common practice is that a plastic change is assumed to be adaptive when its direction is the same as that of the subsequent genetic change or when it moves the trait closer to the state represented by the population adapted to the new environment [29, 30], and we have followed this practice in our analysis.

Another caveat is that the expression levels of different genes may not be independent from one another in the sense that they could be affected by the same mutations, but the same can be said of other types of traits. Hence, the focus on gene expression levels in our study does not seem to cause problems that do not exist in past studies of genetic assimilation of other types of traits. Yet, analyzing all genes in a transcriptome offers a systematic and unbiased view of all traits of one type. This said, because the gene expression levels of each multicellular species analyzed in our study mainly came from one time point and a limited number of separate tissues, genetic assimilations of certain expression traits such as tissue/cell-specific or developmental stage-specific gene expressions are unlikely to be detected here. For example, fly sensitivity to the ether treatment in generating the *bithorax* phenotype in Waddington’s experiment is correlated with *Ubx* expression in the third thoracic imaginal disc [53]. Furthermore, *Ubx* up- and down-regulates hundreds of distinct target genes at specific stages of haltere development [54]. Expression data from multiple tissues (or cell types) and developmental stages would be necessary to probe the potential assimilation of such expression traits.

Although our results were obtained exclusively from gene expression traits, the reasoning why genetic assimilation is not expected in environmental adaptations may apply to all types of phenotypic traits. This prediction should be scrutinized by future studies of other phenotypic traits. We found that gene expressions controlled by fewer TFs are more likely than those controlled by more TFs to show genetic assimilation. For other types of traits, it may also be true that traits controlled by fewer genes are more likely than those controlled by more genes to exhibit genetic assimilation. It would be interesting to test this hypothesis in the future. Our results, along with the previous finding that plasticity is more often maladaptive than adaptive [29–32], suggest that the plasticity-first model of environmental adaptation has limited applicability.

## MATERIALS AND METHODS

### *E. coli* transcriptomic data

Horinouchi *et al*. [33] performed experimental evolution of *E. coli* MDS42 cells first in the M9 medium with glucose (5 g/liter) for approximately 90 generations and then in one of 11 different harsh environments for 350 to 800 generations. The 11 conditions used were M9 medium with 400 mM sodium chloride (NaCl), 210 mM potassium chloride (KCl), 16 μM cobalt chloride (CoCl_2_), 32.5 mM sodium carbonate (Na_2_CO_3_), 40 mM L-lactate (Lac), 30 mM L-malate (Mal), 8.75 mM methacrylate (MCL), 50 mM crotonate (Cro), 350 μM methylglyoxal (MG), 1.25% *n*-butanol (BuOH), and 4.8 μM cetylpyridinium chloride (CPC), respectively. This experiment was performed with five replicates per environment. Before RNA extraction, *E. coli* cells were precultured for 60 hrs under the conditions to be tested. Transcriptomic data were collected using microarrays for (i) three replicate transcriptomes of the ancestral strain in M9 (state O), (ii) one transcriptome of the ancestral strain in each of the 11 harsh environments (state P), (iii) one transcriptome of each of the five replicates of a strain adapted to each of the 11 harsh environments in the respective harsh environment (state A), (iv) one transcriptome of each of the five replicates of a strain adapted to each of the 11 harsh environments in M9 (state B). All transcriptomic data were from Table S2 in Horinouchi *et al*. [33].

### Yeast transcriptomic data

The yeast dataset was generated from the experimental evolution of the haploid strain BY4741. Briefly, Dhar *et al*. [34] evolved three replicate populations under salt stress—YPG (2% peptone + 1% yeast extract + 2% galactose) supplemented with 0.5 M NaCl, and three replicate populations under oxidative stress—YPG supplemented with 1 mM H_2_O_2_, each for 300 generations. The following transcriptomic data were collected using microarrays: (i) two replicate transcriptomes of the ancestral strain in YPG (state O), (ii) two replicate transcriptomes of the ancestral strain in salt and oxidative stresses, respectively (state P), (iii) four transcriptomes from the three replicates of salt-evolved populations under salt stress (two replicate transcriptomes from one of the replicates and one transcriptome from each of the other two replicates), and one transcriptome of each of the three replicate oxidative-evolved populations under oxidative stress (state A), and (iv) one transcriptome of each replicate of salt-evolved populations and oxidative-evolved populations in YPG (state B). The expression data were retrieved from Dhar *et al*. [34].

### Morning glory transcriptomic data

Van Etten *et al*. generated the transcriptome data after experimental evolution of the common morning glory [35]. The ancestral population (*n* = 122) was sampled from the University of Georgia Plant Sciences Farm in Oconee, GA in 2000. The offspring of the population were screened in a greenhouse for high resistance to the herbicide glyphosate. The top 20% most resistant lines were used to select for herbicide resistance for three generations. They also randomly chose 20% lines from the ancestral population and evolved them in a nonselected environment for the same number of generations as a control. They then planted seeds from evolved lines and control lines. After six weeks of growth, they sprayed the herbicide in both lines and collected leaf tissues 8 and 32 hrs after spraying. The following leaf transcriptomic data were collected by RNA sequencing (RNA-seq): (i) herbicide-selected lines both 8 and 32 hrs after herbicide application (state A), (ii) selected lines in the herbicide-free environment at the same two time points (state B), (iii) non-selected control lines 8 and 32 hrs after herbicide application (state P), and (iv) control lines in the herbicide-free environment at the same two time points (state O). Each of these four states at each time point had seven to eight replicates.

### Fruit fly transcriptomic data

Mallard *et al*. [36, 37] exposed ten replicates of a fruit fly population to two different environments: a hot environment with 12 hrs at 18°C (dark) and 12 hrs at 28°C (light) per day and a cold environment with 12 hrs at 10°C (dark) and 12 hrs at 20°C (light) per day, for a total of 64 (hot) and 39 (cold) generations, respectively. Transcriptomic data were collected using RNA-seq from 3- to 5-day-old males in four populations: (i) five replicates of cold-evolved populations in the common garden experiment at 15°C (average temperature of the cold environment) (state O); (ii) five replicates of cold-evolved populations in the common garden experiment at 23°C (average temperature of the hot environment) (state P), (iii) five replicates of hot-evolved populations in the common garden experiment at 23°C (state A), and (iv) five replicates of hot-evolved populations in the common garden experiment at 15°C (state B).

### Copepod transcriptomic data

Brennan *et al*. [38] evolved copepods originally collected in 2016 from Esker Point Beach in Groton under a condition representing today’s environment (18°C, pH ~8.2, and pCO2 ~400□μatm), as well as under a condition representing a potential future environment (22°C, pH ~7.5, and pCO2 ~2000□μatm) for 20 generations. At the 21st generation, they reciprocally transplanted some of the copepods and kept them for three generations, in addition to keeping the rest of the copepods under their previously adapted conditions for the same number of generations. Each of these four treatments had four replicates. RNA-seq was performed for each replicate of each treatment at the 22nd, 23rd, and 24th generations. We analyzed the data collected at the 24th generation to ensure that the potential plasticity is fully induced upon the transplant.

### Guppy transcriptomic data

Ghalambor and colleagues [29] conducted experimental evolution of Trinidadian guppies that had been adapted to a high-predation environment in two different low-predation environments. After one year of experimental evolution (~3 to 4 generations), fish were collected from each introduced population (Intro1 and Intro2). In addition, they collected fish that had originated from a high-predation environment and were naturally adapted to a low-predation environment. Breeds of the ancestral population sampled from the high-predation environment were respectively reared in a low-predation environment (state P) and a high-predation environment (state O) in the laboratory. Breeds of Intro1, Intro2, and the naturally adapted low-predation population were also respectively reared in the low- (state A) and high-predation (state B) environments. Brain transcriptomes were assessed by RNA-seq. Numbers of fish individuals used were five for the ancestral high-predation population reared in each environment, four for the natural low-predation population reared in the high-predation environment, two for the natural low-predation population reared in the low-predation environment, and four for Intro1 and Intro2 reared in each environment. Note that the original authors reported concordantly differentially expressed genes across experiments [29], which differ from DEGs of each experiment analyzed here.

### Chicken transcriptomic data

Ho *et al*. [39] performed RNA-seq of five tissues (brain, heart, liver, lung, and muscle) from Tibetan chickens raised in A’ba (3300 m above sea level) and lowland chickens raised in Ya’an (670 m above sea level). These two locations are on the opposite sides of the east edge of the Tibet Plateau in Sichuan Province, China. They also performed reciprocal transplant experiments (hatched and raised lowland chickens in the highland and highland chickens in the lowland), followed by transcriptome profiling of the same five tissues. For each of the four groups of chickens, transcriptomes were collected from healthy males of similar body weights at the age of 120 days, including eight to nine chickens for each tissue except for the muscle, which had five to six chickens.

### Analysis of RNA-seq data

Gene expressions were measured by RNA-seq in all studies analyzed except for *E. coli* and yeast. We obtained the read counts of each gene archived by the original authors. DESeq2 [55] was used to identify differentially expressed genes (DEGs) between *E*_O_ and *E*_A_ using an FDR of 0.05. A DEG is considered a PSG if *E*_P_ is significantly different from *E*_O_ but not significantly different from *E*_A_ based on DESeq2 (FDR = 0.05 in each test, considering all DEGs). A PSG is considered to support genetic assimilation if *E*_B_ is significantly different from *E*_O_ but not significantly different from *E*_A_ based on DESeq2 (FDR = 0.05 in each test, considering all PSGs). Similarly, a PSG is considered to refute genetic assimilation if *E*_B_ is significantly different from *E*_A_ but not significantly different from *E*_O_ based on DESeq2 (FDR = 0.05 in each test, considering all PSGs). PGGs/APGs and the subset of PGGs/APGs supporting or refuting genetic assimilation were similarly identified.

### Analysis of microarray data

Gene expressions were measured using microarrays in the studies of *E. coli* [33] and yeast [34]. The background-corrected intensity values were normalized using the quantile normalization method [56] in both studies. We obtained the gene expression levels archived by the original authors. We identified DEGs, PSGs (or PGGs/APGs), and PSGs (or PGGs/APGs) that support or refute genetic assimilation as described in the preceding section, except that the R package limma [57] instead of DESeq2 was used.

### Gene Ontology (GO) analysis

GO annotations of all organisms studied were downloaded from the Gene Ontology resource (http://geneontology.org) [58]. GO enrichment for genes supporting genetic assimilation was calculated in R using the topGO v2.44.0 package [59] with the following parameters (ontology = “BP/MF/CC”, testStatistic = GOFisherTest, and classic Fisher test *P* < 0.01). We used the corresponding PSGs as the background set of genes in the enrichment test. Following the instructions of the package, we report *P* values from the weight algorithm without multiple testing correction.

### Transcription factors (TFs)

The number of TFs controlling each *E. coli* gene was obtained from RegulonDB v10.5 [60]. Specifically, RegulonDB lists 210 TFs that each control at least one target gene. In total, these 210 TFs control 1,864 genes via 4,519 TF-target gene interactions.

## ACKNOWLEDGEMENTS

We are grateful to the authors of the original studies for sharing the gene expression data and members of the Zhang laboratory for valuable comments.

## FIGURE LEGENDS

### Legends of supplementary figures

**Fig. S1. Genes supporting or refuting genetic assimilation in laboratory adaptations of *E. coli* to 11 different stressful environments.** Same as Fig. 2a except that the same sample size of three replicates is used in *E*_A_ and *E*_O_ measurement.

**Fig. S2. Number of DEGs between the ancestral population and cold-evolved population and number of DEGs between the ancestral population and hot-evolved population. a**, Comparison between the two DEG numbers when gene expression is measured in the cold environment (15°C). **b**, Comparison between the two DEG numbers when gene expression is measured in the hot environment (23°C). *P*-values are based on two-tailed binomial tests of equality between the two bars.

**Fig. S3. GO enrichment of *E. coli* genes exhibiting genetic assimilation in at least one of the 11 environments, relative to all PSGs.**

**Fig. S4. Genes supporting or refuting genetic assimilation in laboratory adaptations of *E. coli* to 11 different stressful environments.** Same as Fig. 2a, except that adaptive plasticity genes (APGs) instead of PSGs are examined.

**Fig. S5. Genes supporting or refuting genetic assimilation in five other species studied.** Same as Fig. 3, except that APGs instead of PSGs are examined.

**Fig. S6. Genes supporting or refuting genetic assimilation in six species studied.** Same as Fig. 2a and Fig. 3, except that PGGs instead of PSGs are examined.

**Fig. S7. Numbers of genes that gained or lost plasticity upon environmental adaptations in the seven species studied.** Group 1 genes (brown bar) are significantly plastic in the ancestral genotype but not significantly plastic in the derived genotype, whereas Group 2 genes (green bar) are significantly plastic in the derived genotype but not significantly plastic in the ancestral genotype. In *E. coli*, yeast, guppy, and chicken, the combined results from all environments or tissues examined are shown. *P*-values are from two-tailed binomial tests of equality between Group 1 and Group 2 gene numbers.

## Notes

### Competing Interest Statement

The authors have declared no competing interest.

