## Supplementary Figures 1-7 for "Transcriptomic analysis of diverse organisms reveals the rareness of genetic assimilation in environmental adaptations"

**This PDF includes:**

Legends of supplementary figures

Figures S1 to S7

**Legends of supplementary figures**

**Fig. S1. Genes supporting or refuting genetic assimilation in laboratory adaptations of *E. coli* to 11 different stressful environments.** Same as Fig. 2a except that the same sample size of three replicates is used in $E_A$ and $E_O$ measurement.

**Fig. S2. Number of DEGs between the ancestral population and cold-evolved population and number of DEGs between the ancestral population and hot-evolved population. a**, Comparison between the two DEG numbers when gene expression is measured in the cold environment (15°C). **b**, Comparison between the two DEG numbers when gene expression is measured in the hot environment (23°C). *P*-values are based on two-tailed binomial tests of equality between the two bars.

**Fig. S3. GO enrichment of *E. coli* genes exhibiting genetic assimilation in at least one of the 11 environments, relative to all PSGs.**

**Fig. S4. Genes supporting or refuting genetic assimilation in laboratory adaptations of *E. coli* to 11 different stressful environments.** Same as Fig. 2a, except that adaptive plasticity genes (APGs) instead of PSGs are examined.

**Fig. S5. Genes supporting or refuting genetic assimilation in five other species studied.** Same as Fig. 3, except that APGs instead of PSGs are examined.

**Fig. S6. Genes supporting or refuting genetic assimilation in six species studied.** Same as Fig. 2a and Fig. 3, except that PGGs instead of PSGs are examined.

**Fig. S7. Numbers of genes that gained or lost plasticity upon environmental adaptations in the seven species studied.** Group 1 genes (brown bar) are significantly plastic in the ancestral genotype but not significantly plastic in the derived genotype, whereas Group 2 genes (green bar) are significantly plastic in the derived genotype but not significantly plastic in the ancestral genotype. In *E. coli*, yeast, guppy, and chicken, the combined results from all environments or tissues examined are shown. *P*-values are from two-tailed binomial tests of equality between Group 1 and Group 2 gene numbers.

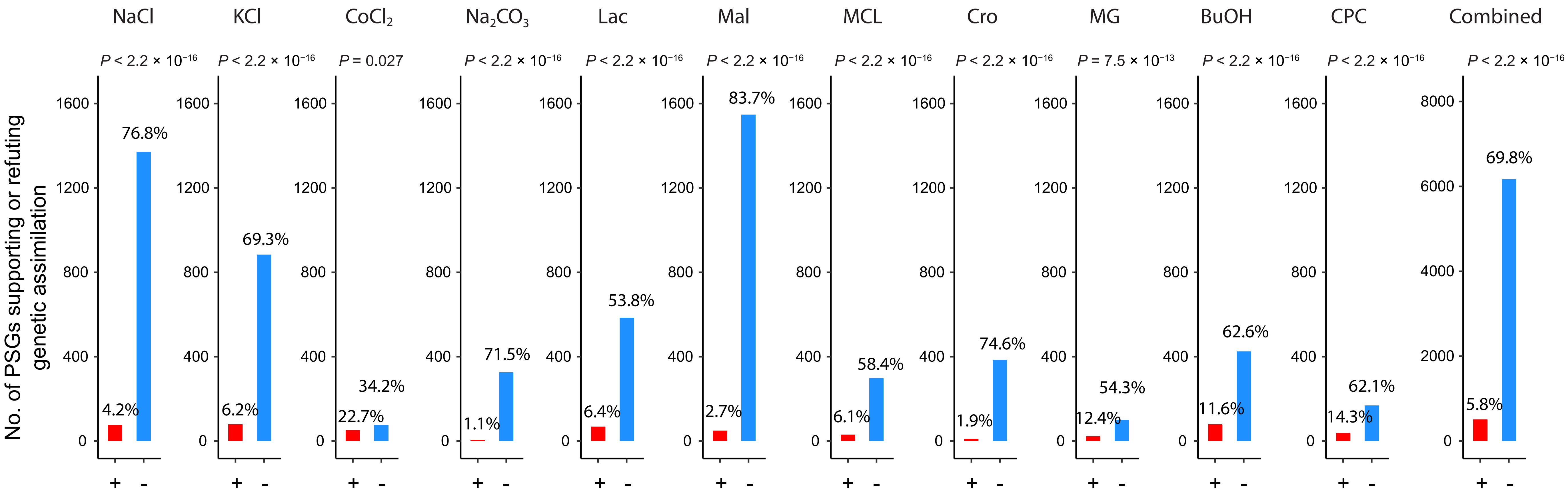

No. of PSGs supporting or refuting genetic assimilation
NaCl
KCl
$CoCl_2$
$Na_2CO_3$
Lac
Mal
MCL
Cro
MG
BuOH
CPC
Combined
$P < 2.2 \times 10^{-16}$
$P < 2.2 \times 10^{-16}$
$P = 0.027$
$P < 2.2 \times 10^{-16}$
$P < 2.2 \times 10^{-16}$
$P < 2.2 \times 10^{-16}$
$P < 2.2 \times 10^{-16}$
$P < 2.2 \times 10^{-16}$
$P = 7.5 \times 10^{-13}$
$P < 2.2 \times 10^{-16}$
$P < 2.2 \times 10^{-16}$
$P < 2.2 \times 10^{-16}$
4.2%
76.8%
6.2%
69.3%
22.7%
34.2%
1.1%
71.5%
6.4%
53.8%
2.7%
83.7%
6.1%
58.4%
1.9%
74.6%
12.4%
54.3%
11.6%
62.6%
14.3%
62.1%
5.8%
69.8%
0
400
800
1200
1600
2000
4000
6000
8000
+
-


Fig. S1

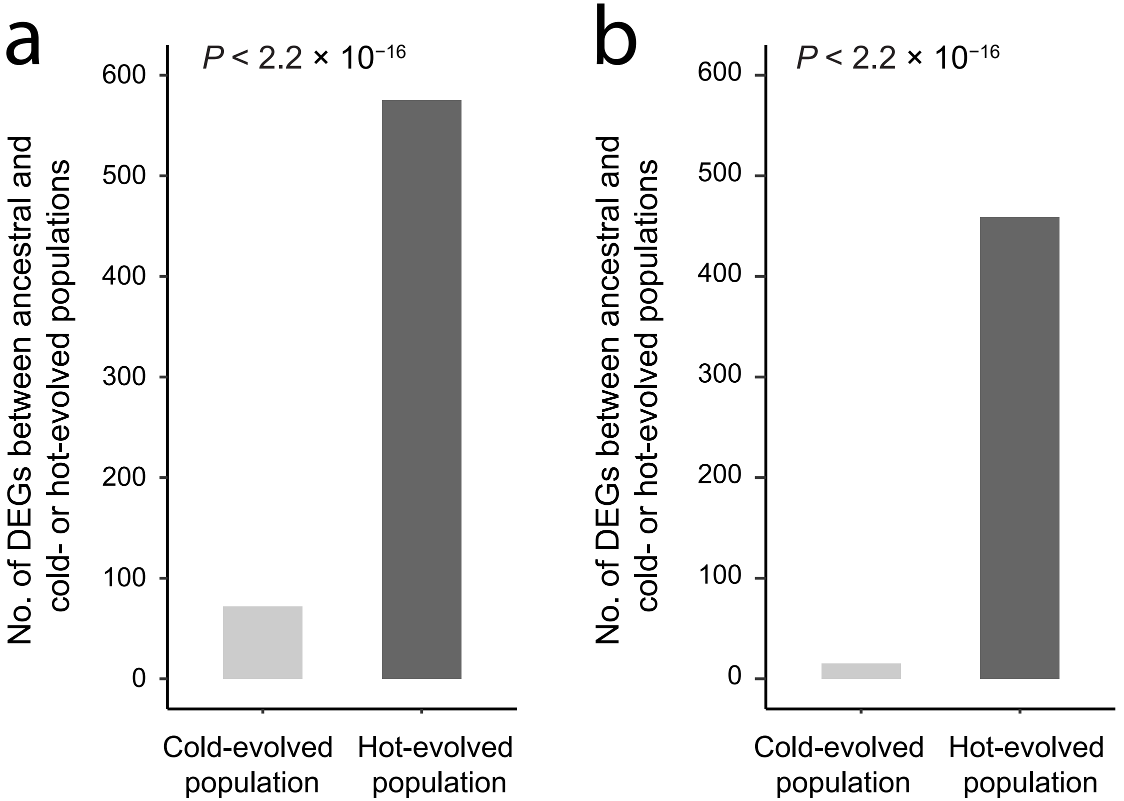
a
P < 2.2 × 10−16
No. of DEGs between ancestral and cold- or hot-evolved populations
0
100
200
300
400
500
600
Cold-evolved population
Hot-evolved population
b
P < 2.2 × 10−16
No. of DEGs between ancestral and cold- or hot-evolved populations
0
100
200
300
400
500
600
Cold-evolved population
Hot-evolved population

Fig. S2

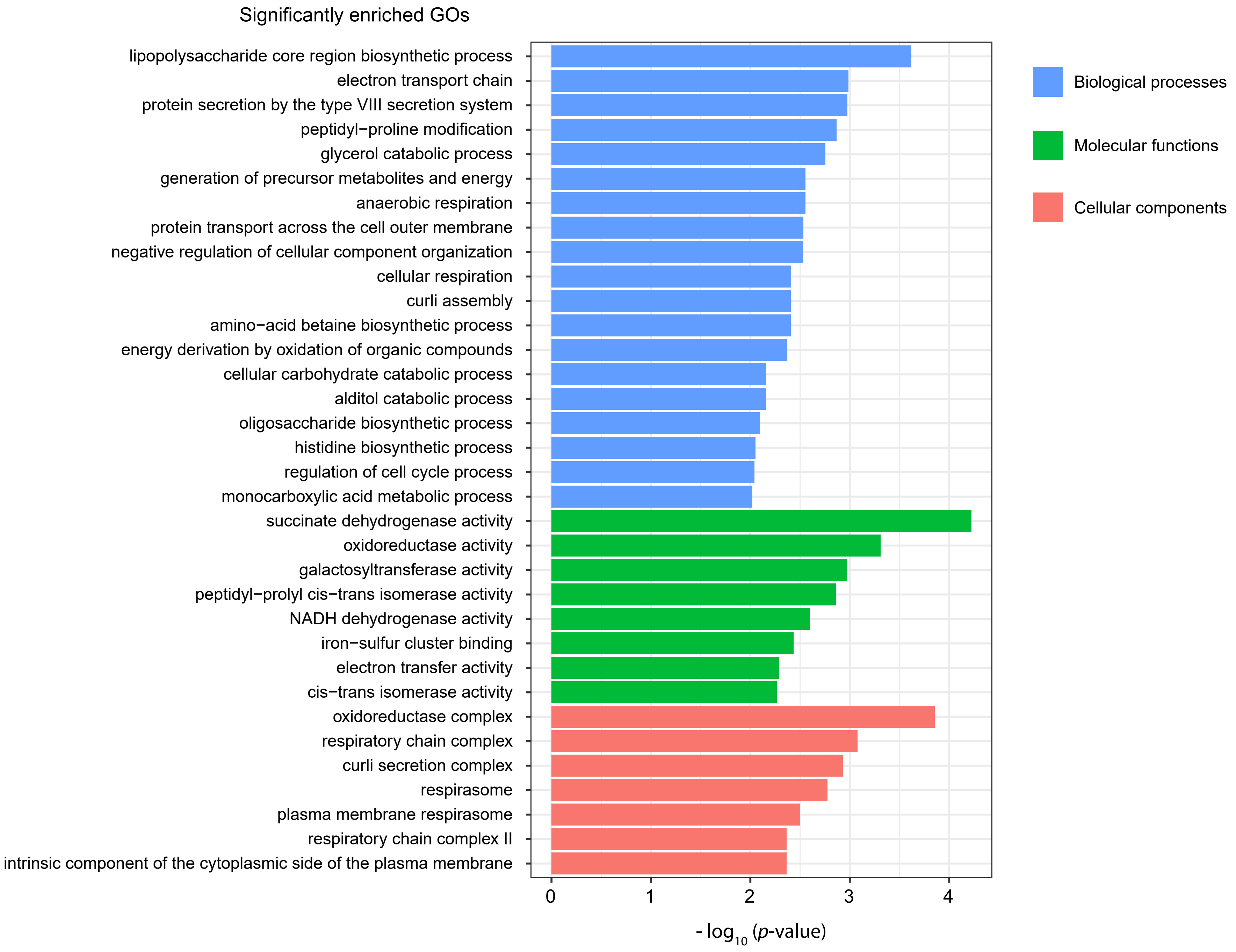

Significantly enriched GOs
lipopolysaccharide core region biosynthetic process
electron transport chain
protein secretion by the type VIII secretion system
peptidyl−proline modification
glycerol catabolic process
generation of precursor metabolites and energy
anaerobic respiration
protein transport across the cell outer membrane
negative regulation of cellular component organization
cellular respiration
curli assembly
amino−acid betaine biosynthetic process
energy derivation by oxidation of organic compounds
cellular carbohydrate catabolic process
alditol catabolic process
oligosaccharide biosynthetic process
histidine biosynthetic process
regulation of cell cycle process
monocarboxylic acid metabolic process
succinate dehydrogenase activity
oxidoreductase activity
galactosyltransferase activity
peptidyl−prolyl cis−trans isomerase activity
NADH dehydrogenase activity
iron−sulfur cluster binding
electron transfer activity
cis−trans isomerase activity
oxidoreductase complex
respiratory chain complex
curli secretion complex
respirasome
plasma membrane respirasome
respiratory chain complex II
intrinsic component of the cytoplasmic side of the plasma membrane
0
1
2
3
4
- log10 (p-value)
Biological processes
Molecular functions
Cellular components


Fig. S3

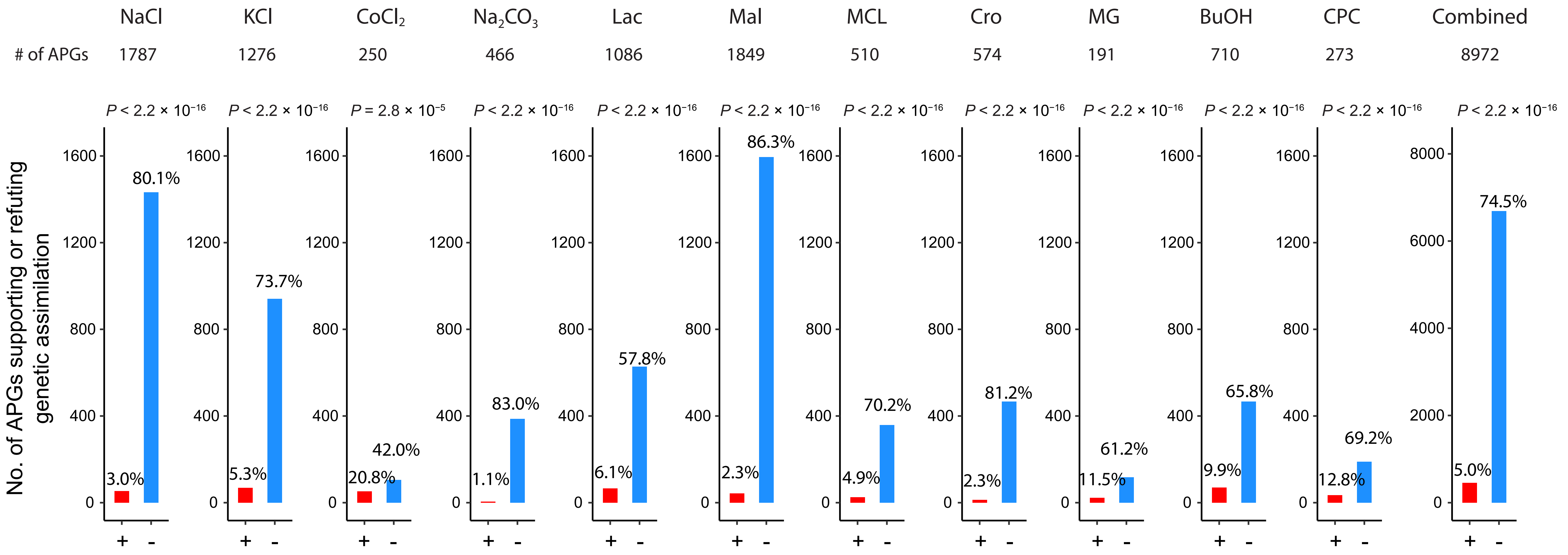
NaCl
KCl
$CoCl_2$
$Na_2CO_3$
Lac
Mal
MCL
Cro
MG
BuOH
CPC
Combined
### of APGs
1787
1276
250
466
1086
1849
510
574
191
710
273
8972
$P < 2.2 \times 10^{-16}$
$P = 2.8 \times 10^{-5}$
No. of APGs supporting or refuting genetic assimilation
3.0%
80.1%
5.3%
73.7%
20.8%
42.0%
1.1%
83.0%
6.1%
57.8%
2.3%
86.3%
4.9%
70.2%
2.3%
81.2%
11.5%
61.2%
9.9%
65.8%
12.8%
69.2%
5.0%
74.5%
+
-

Fig. S4

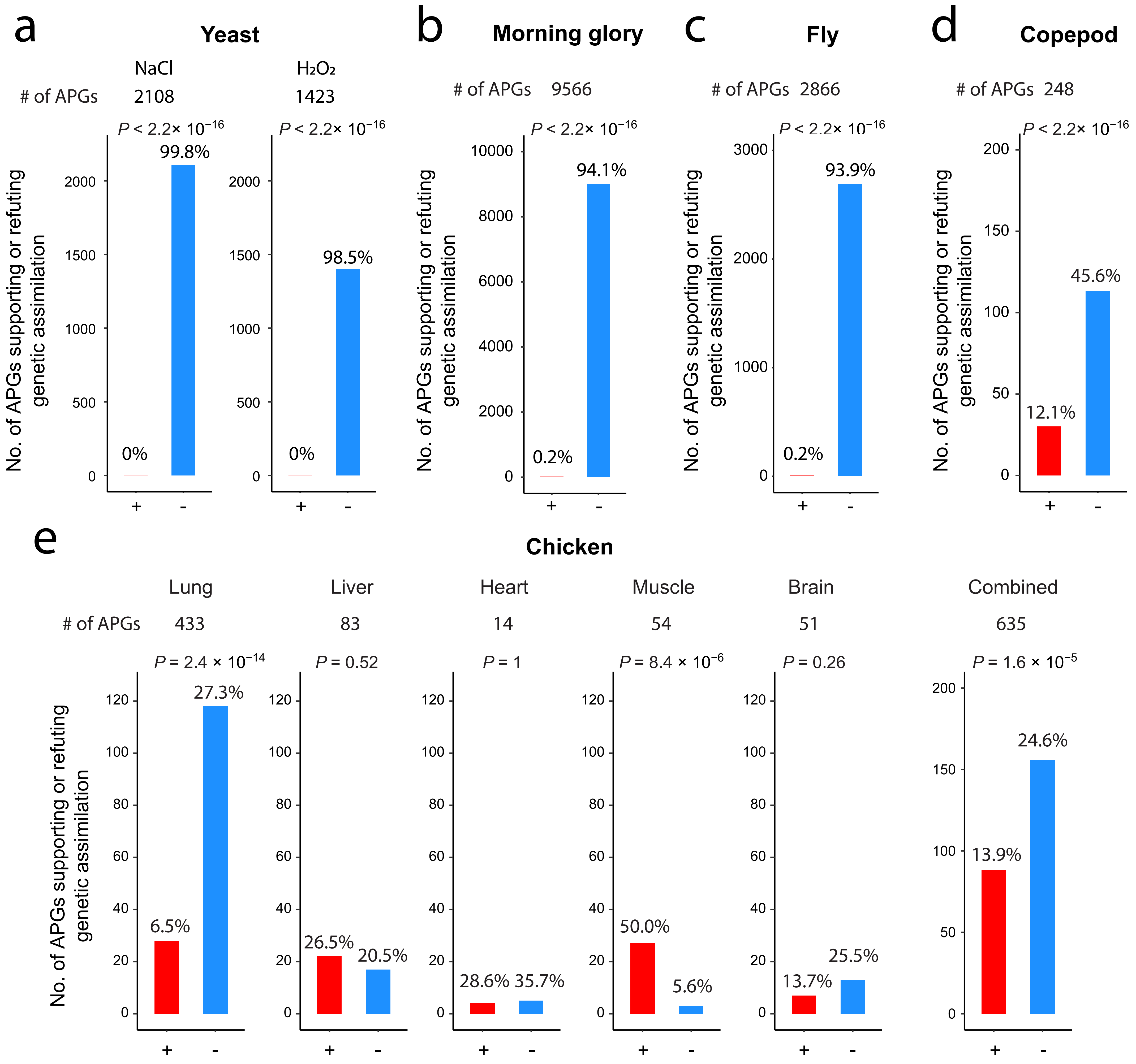

a
Yeast
NaCl
H2O2
### of APGs 2108 1423
P < 2.2× 10−16
99.8%
0%
98.5%
0%
b
Morning glory
### of APGs 9566
P < 2.2× 10−16
94.1%
0.2%
c
Fly
### of APGs 2866
P < 2.2× 10−16
93.9%
0.2%
d
Copepod
### of APGs 248
P < 2.2× 10−16
45.6%
12.1%
No. of APGs supporting or refuting genetic assimilation
+
-
e
Chicken
Lung Liver Heart Muscle Brain Combined
### of APGs 433 83 14 54 51 635
P = 2.4 × 10−14
6.5%
27.3%
P = 0.52
26.5%
20.5%
P = 1
28.6% 35.7%
P = 8.4 × 10−6
50.0%
5.6%
P = 0.26
13.7%
25.5%
P = 1.6 × 10−5
13.9%
24.6%


Fig. S5

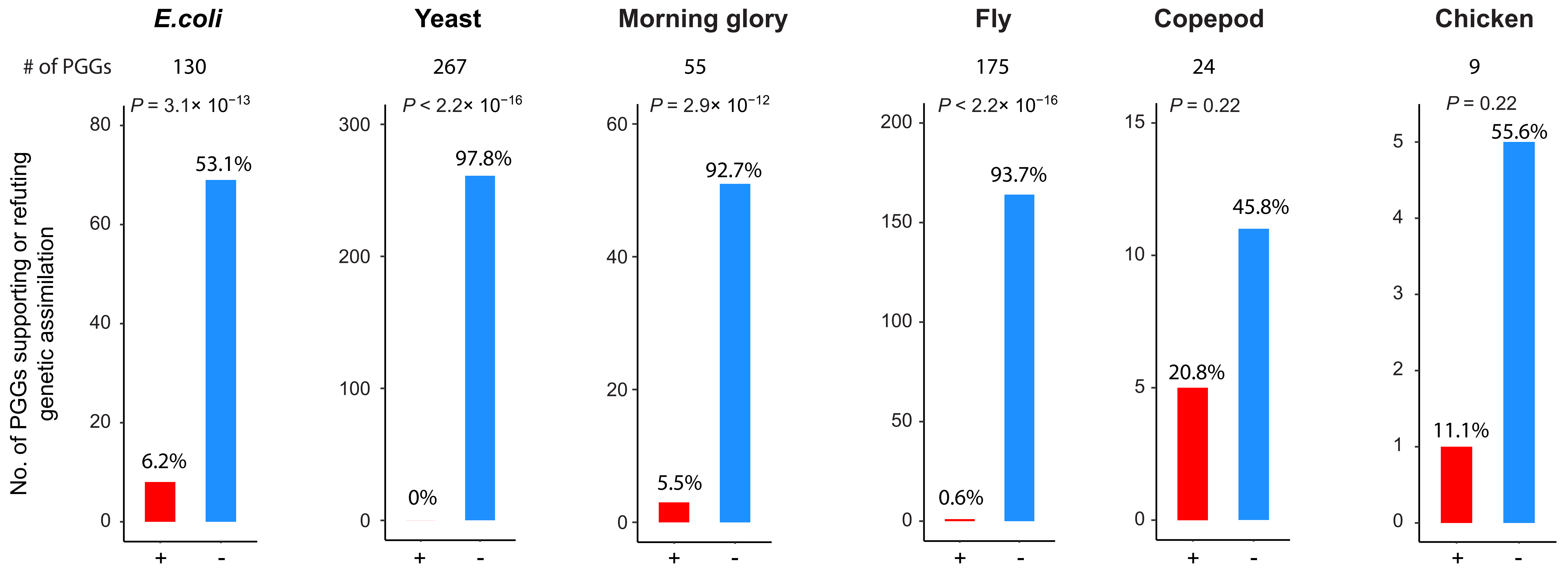
E.coli
Yeast
Morning glory
Fly
Copepod
Chicken
### of PGGs
130
267
55
175
24
9
P = 3.1× 10^−13
P < 2.2× 10^−16
P = 2.9× 10^−12
P < 2.2× 10^−16
P = 0.22
P = 0.22
No. of PGGs supporting or refuting genetic assimilation
6.2%
53.1%
0%
97.8%
5.5%
92.7%
0.6%
93.7%
20.8%
45.8%
11.1%
55.6%
+
-

Fig. S6

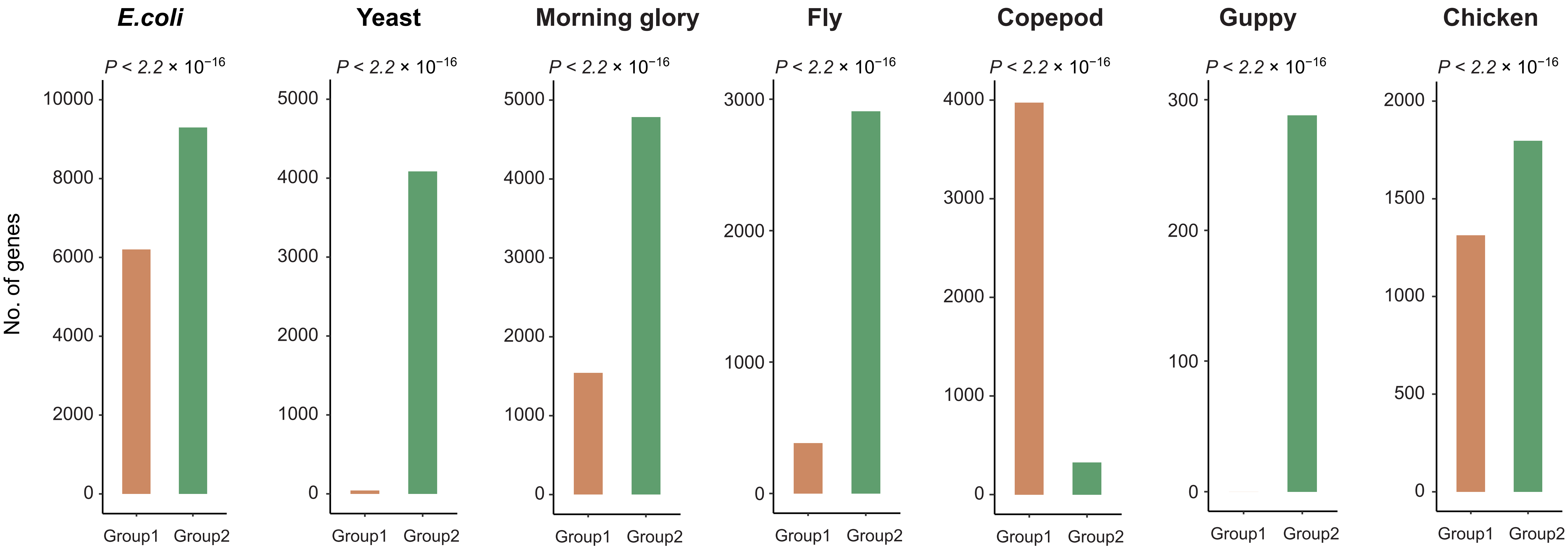
E.coli
Yeast
Morning glory
Fly
Copepod
Guppy
Chicken
P < 2.2 × 10−16
No. of genes
Group1
Group2
10000
8000
6000
4000
2000
0
5000
4000
3000
2000
1000
3000
300
200
100
1500
500

Fig. S7
